# One score to rule them all: severity assessment in laboratory mice

**DOI:** 10.1101/2020.06.23.166801

**Authors:** Steven R. Talbot, Birgitta Struve, Laura Wassermann, Miriam Heider, Nora Weegh, Tilo Knape, Martine C. J. Hofmann, Andreas von Knethen, Paulin Jirkof, Lydia Keubler, André Bleich, Christine Häger

## Abstract

Animal welfare and the refinement of experimental procedures are fundamental aspects of biomedical research. They provide the basis for robust experimental designs and reproducibility of results. In many countries, the determination of welfare is a mandatory legal requirement and implies the assessment of the degree of the severity that an animal experiences during an experiment. However, for an effective severity assessment, an objective and exact approach/system/strategy is needed. In light of these demands, we have developed the Relative Severity Assessment (RELSA) score.

This comprehensive composite score was established on the basis of physiological and behavioral data from a surgical mouse study. Body weight, the Mouse Grimace Scale score, burrowing behavior, and the telemetry-derived parameters heart rate, heart rate variability, temperature, and general activity were used to investigate the quality of indicating severity during postoperative recovery. The RELSA scores not only revealed individual severity levels but also allowed a comparison of severity in distinct mouse models addressing colitis, sepsis, and restraint stress using a *k*-means clustering approach with the maximum achieved RELSA scores.

We discriminated and classified data from sepsis nonsurvivors into the highest relative severity level. Data from mice after intraperitoneal transmitter implantation and sepsis survivor al were located in the next lower cluster, while data from mice subjected to colitis and restraint stress were placed in the lowest severity cluster. Analysis of individual variables and their combinations revealed model- and time-dependent contributions to severity levels.

In conclusion, we propose the RELSA score as a validated tool for objective real-time applicability in severity assessment and as a first step towards a unified and accessible risk assessment tool in biomedical research. As an effective severity assessment system, it will fundamentally improve animal welfare, as well as data quality and reproducibility.

## Introduction

Good science and high quality data derived from animal experiments in basic and translational research requires good animal welfare. Consequently, researchers are obligated to ensure the best possible welfare of research animals, in line with the refinement principle in the 3Rs^1,2^. Therefore, determination of the welfare of animals under scientific procedures is embedded in many international animal protection guidelines and acts, e.g., the Guide for the Care and Use of Laboratory Animals^3^ or the European Directive on the protection of animals used for scientific purposes^4^. Animal welfare describes a status of high quality of life, which relies on the consideration and promotion of things to achieve good animal welfare^5,6^. Its assessment requires the monitoring of animal affective states with both positive and negative valance^7^. The term severity assessment more specifically addresses the categorization of negative affective states of animals under experimental conditions; however, both the promotion of welfare and assessment of severity aim at the recognition of any sign of suffering and thereby are essential for possible interventions to relieve the burden during experiments and promote the refinement of procedures. Nevertheless, to achieve compliance with scientific and regulatory requirements, an exact, evidence-based severity assessment and the classification of severity conditions are needed^8^.

Evidence-based severity assessment also provides the basis for disclosing information on animal welfare in scientific studies. According to the ARRIVE guidelines, welfare conditions related to animal housing, handling, analgesia, and other procedures should be made transparent to the research community^9^. This openness is required not only to overcome the reproducibility crisis^10,11^ but also to advance the whole field/subject to a more objective level of comparability in terms of results, in this case severity assessment. Prominent methods for assessing the severity of the animal’s burden include the measurement of multiple variables coming from various scientific areas, resulting in a highly interdisciplinary and growing field of research. Currently, parameters from the field of physiology, biochemistry, clinical sciences and behavioral sciences are considered to best reflect the welfare state of the animals under experimentation^7^. While there are many potential sources of information and methods to assess severity on a single or multiparameter basis, there has been almost no attempt to a) algorithmically find the best indicator or set of indicators for a specific question, e.g., to assess the impact of a specific scientific procedure or model system on the animal or to differentiate short or long-lasting impacts on welfare, or b) build data-driven composite metrics for variables coming from different sources. This leads to a situation in which variable-bound criteria (e.g., a loss of body weight) are frequently combined with subjective clinical scoring methods to assess animal welfare states, which are then used for setting “humane endpoints”^12^.

To obtain a clearer, more precise and holistic picture of impaired animals, multimodal and multivariate methods need to be integrated into severity assessment. This integration requires adequate compositional scoring systems. We have previously contributed evidence that the inclusion of data science and abstract mathematical methods to explain higher-order relationships in the data that go beyond the traditional scope of statistical *post hoc* analyses is highly informative in terms of determining severity conditions or levels^13–15^. Building on this experience, we aimed to develop an algorithm-based composite scoring system that enables objective monitoring of welfare states using a theoretically indefinite number of variables from various scientific fields, allowing comparisons between individual animals, treatment groups and animal models. This system should also be able to depict the best parameters or their combinations to assess the animals.

In this study, a broad set of physiological, clinical and behavioral data from mouse models involving surgery and pain, inflammation, stress, and sepsis with varying degrees of severity were used to develop and validate the Relative Severity Assessment (RELSA) score using a common scale for multivariate input data. Importantly, the algorithm-based score enables real-time severity quantification. Additionally, it not only reveals severity of an individual but also enables the comparison of animal models using a *k*-means clustering approach. Therefore, the RELSA score developed in this study is model independent and can be established with random variables to characterize the severity of any condition with adverse effects, possibly even in human patients in the near future.

## Results

### The quality of physiological and behavioral variables for the characterization of welfare states

First, we monitored the impact of intraperitoneal transmitter implantation (TM) or corresponding sham surgery on the well-being of mice treated with metamizole or either carprofen for analgesia using distinct behavioral (activity [act], burrowing [bur], Mouse Grimace Scale [mgs]) and physiological variables (body weight change [bwc], temperature [temp], heart rate [hr], heart rate variability [hrv]). The measured variables showed their most extreme changes immediately after surgery (post-op) and over the subsequent 4 days. While we found significant differences between TM-implanted and sham-operated animals, we did not detect relevant differences between the utilized analgesic regimens carprofen and metamizole (Fig. S1a-h, S2a-d). For an overall impression of the variable changes, the Cohen’s d effect sizes were determined for all TM-implanted and sham-operated animals. Depending on the nature of each measured variable, either a drop or an increase in values was observed compared to baseline (bsl), thereby reflecting effects on animal welfare in a time-dependent manner (Fig. 1). Integrative principal component analysis (PCA) with data on post-op day, day 1, day 2, day 4 and day 7, as well as physiological values, revealed no strong dominant variable in the first principal component for the TM group (Fig. 2a). Here, variable contributions were slightly higher than the uniform distribution of variance in all factors (12.5%; red dashed line, Fig. 2b). The second-largest contribution, as shown in PC2, was dominated by temperature (51.98%). The two-dimensional data showed a time-dependent flow of clusters starting at baseline values, increases during the post-op day and then slowly back towards physiological normal values. In the sham-operated group, this time-dependent flow was less apparent. Here, an approximate discrimination of the 95% confidence ellipses between the baseline and post-op day was possible. All other days did not allow explicit discrimination (Fig. 2c). Again, no strong variable was identified in PC1. PC2 was dominated by mgs (Fig. 2d).

**Figure 1.**
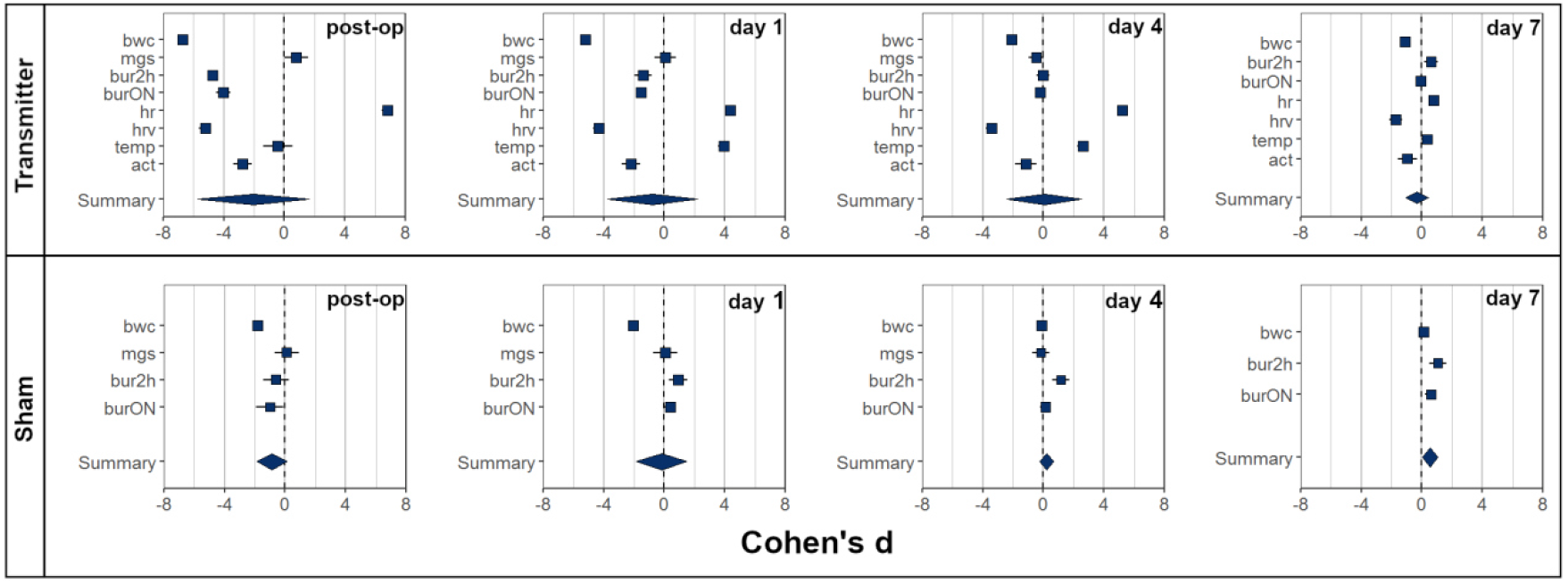
Effect sizes of behavioral and physiological variables after surgical interventions. Effect sizes (ranging from −8 to +8 depending on an increase or decrease in the respective variable) were individually calculated for the variables body weight change (bwc), Mouse Grimace Scale (mgs), burrowing behavior (bur: 2h= start before dark phase, ON= overnight), heart rate (hr), heart rate variability (hrv), body temperature (temp), general activity (act) and summarized (summary of variables) as scaled differences to baseline values using Cohen’s d on the day of surgery (post-op) and on days 1, 4 and 7 after the surgery. In the transmitter- and sham-surgery groups, the effect sizes decrease over time with smaller effect sizes in the sham-operated group. Error bars are 95% confidence intervals.

**Figure 2.**
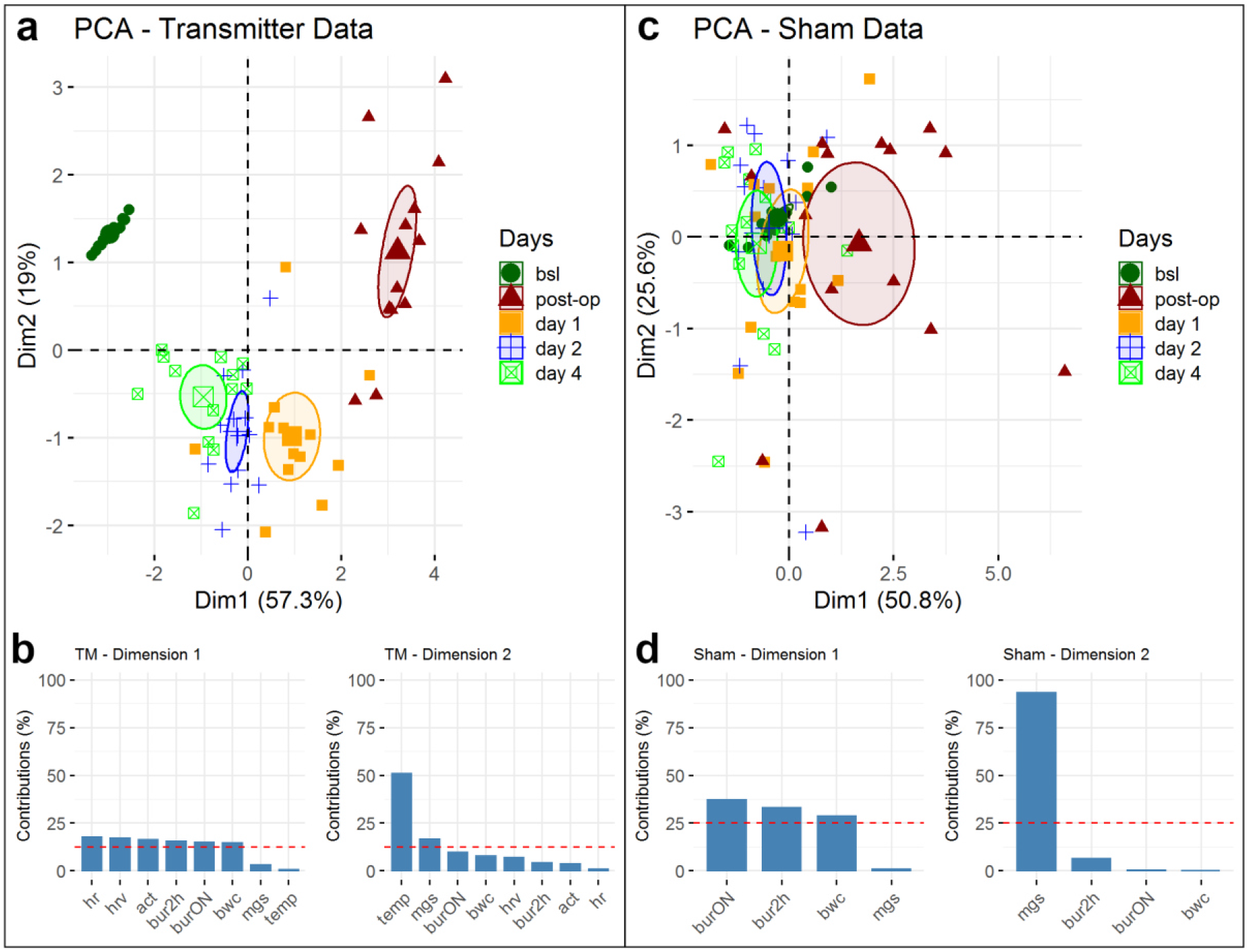
Principal component analysis (PCA) reveals distinct severity states during recovery in a two-dimensional factor space. (**a**) PCA of TM-implanted mice using the variables body weight change (bwc), Mouse Grimace Scale (mgs), burrowing behavior (bur: 2h= start before dark phase, ON= overnight), heart rate (hr), heart rate variability (hrv), body temperature (temp), general activity (act) for baseline, and from postoperative days 1, 2 and 4 after the surgery. The 95% confidence ellipses show good separation on each day while maintaining high total variance in two dimensions (76.3%). (**b**) The strongest dominating variable in dimension 1 is hr with 17.79%, which is marginally above the uniform distribution of 12.5%, and in orthogonal dimension 2, it is temp with 50.85%. (**c**) PCA with sham-operated mice using the variables body weight change (bwc), Mouse Grimace Scale (mgs), burrowing behavior (bur: 2h= start before dark phase, ON= overnight) for baseline and from postoperative days 1, 2 and 4 after the surgery. Here, only the post-op day shows an independent class with a small overlap with the ellipsis of day 1. The remaining days do not show daily separations. The total variance remains high at 76.4%. (**d**) In dimension 1, burON (37.11%) is the strongest contributor, and in dimension 2, it is the mgs (93.32%).

The Cohen’s d and PCA results clearly showed that the measured variables reflected the impact of surgery on animal welfare states in a time-dependent manner. Furthermore, the parameters allowed differentiation of transmitter vs sham operations.

### The chosen variables and their value ranges define the severity continuum and thereby the RELSA variable space

To determine the impact of the given procedures on welfare, not only on qualitative scales but also on quantitative scales, the whole spectrum of each variable needs to be assessed. Table 1 shows the normalized minimum/maximum ranges (in percent) for each of the experiments, treatments and analgesic regimens in the reference study. The normalized differences code for the maximum-experienced quantitative severity in the given data and were used in the definition of a RELSA reference set. Depending on the direction that the variables under duress took (increase/decrease), the minimum or maximum values in the reference set were taken to calculate relative effects for each variable.

**Table 1.** Ranges for all subgroups in the reference model.

| Transmitter + carprofen (%) |  |  |  | Sham + carprofen (%) |  |  |
| --- | --- | --- | --- | --- | --- | --- |
|  | min | max | range | min | max | range |
| bwc | 84.55 | 115.76 | 31.22 | 89.36 | 114.51 | 25.15 |
| bur2h | 0.00 | 234.87 | 234.87 | 37.07 | 477.50 | 440.42 |
| burON | 0.00 | 137.62 | 137.62 | 50.26 | 108.31 | 58.05 |
| hr | 97.68 | 153.39 | 55.71 |  |  |  |
| hrv | 8.09 | 113.04 | 104.95 |  |  |  |
| temp | 97.25 | 103.37 | 6.11 |  |  |  |
| act | 4.34 | 164.65 | 160.31 |  |  |  |
| mgs | 19.05 | 405.88 | 386.83 | 0.00 | 341.67 | 341.67 |

| Transmitter + metamizole (%) |  |  |  | Sham + metamizole (%) |  |  |
| --- | --- | --- | --- | --- | --- | --- |
|  | min | max | range | min | max | range |
| bwc | 86.27 | 115.27 | 29.00 | 90.10 | 117.30 | 27.20 |
| bur2h | 0.00 | 181.40 | 181.40 | 6.43 | 218.66 | 212.22 |
| burON | 0.00 | 119.90 | 119.90 | 73.47 | 112.30 | 38.83 |
| hr | 88.61 | 147.20 | 58.59 |  |  |  |
| hrv | 23.71 | 123.51 | 99.79 |  |  |  |
| temp | 95.80 | 104.41 | 8.61 |  |  |  |
| act | 5.77 | 132.24 | 126.47 |  |  |  |
| mgs | 9.41 | 300.00 | 290.59 | 0.00 | 206.38 | 206.38 |

The analgesic regimes for the TM groups were pooled for the following step (n=13) using eight (TM) or 4 (Sham) input variables.

### Defining the RELSA algorithm

The reference set enabled relating individual animals or groups to the maximum level of severity experienced on a quantitative scale. However, the most important step was the combination of all available variables into a single composite score. We, therefore, generated matrices of standardized differences to weigh each variable’s contributions as a means for obtaining a measure for relative severity grades (Fig. 3), which is described in the Methods section in detail.

**Figure 3.**
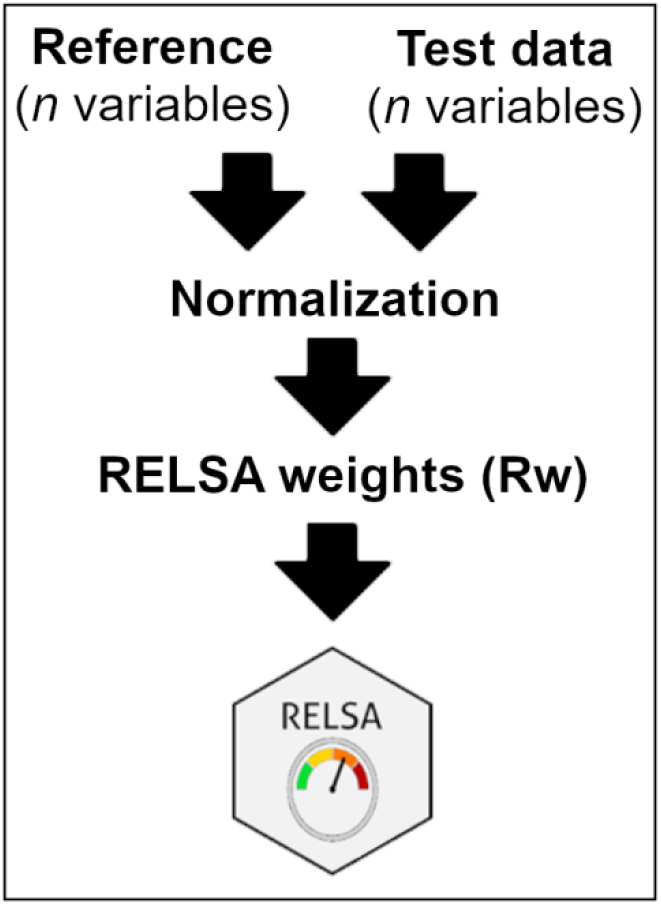
The RELSA workflow. The RELSA algorithm requires a reference data set with *n* variables. Test data require the same variables that are included in the reference set. Both data sources are normalized to their baseline values, followed by the calculation of the individual RELSA weights (R_w_) as standardized effect sizes with regard to the maximal observed changes in the reference set. The final RELSA score is calculated as a root mean square from the available R_w_. The RELSA score can be calculated for single values and for multiple time points.

### RELSA reveals impaired welfare after surgery

RELSA scores were calculated daily for all animals in this study. The averaged scores per day provide an overview of the time course in each of the experimental groups, which were separated into analgesia subgroups for this analysis (Fig. 4a carprofen; Fig. 4b metamizole). Maximum RELSA scores were observed directly post surgery, with mean scores in the TM mice of 0.77 (SD 0.07) for carprofen and 0.73 (SD 0.03) for metamizole. In the sham-operated animals, the mean of the maximum RELSA scores was 0.34 (SD 0.16) for carprofen and 0.44 (SD 0.13) for metamizole, which were also reached directly after surgery. RELSA scores that have returned to zero indicate recovery.

**Figure 4.**
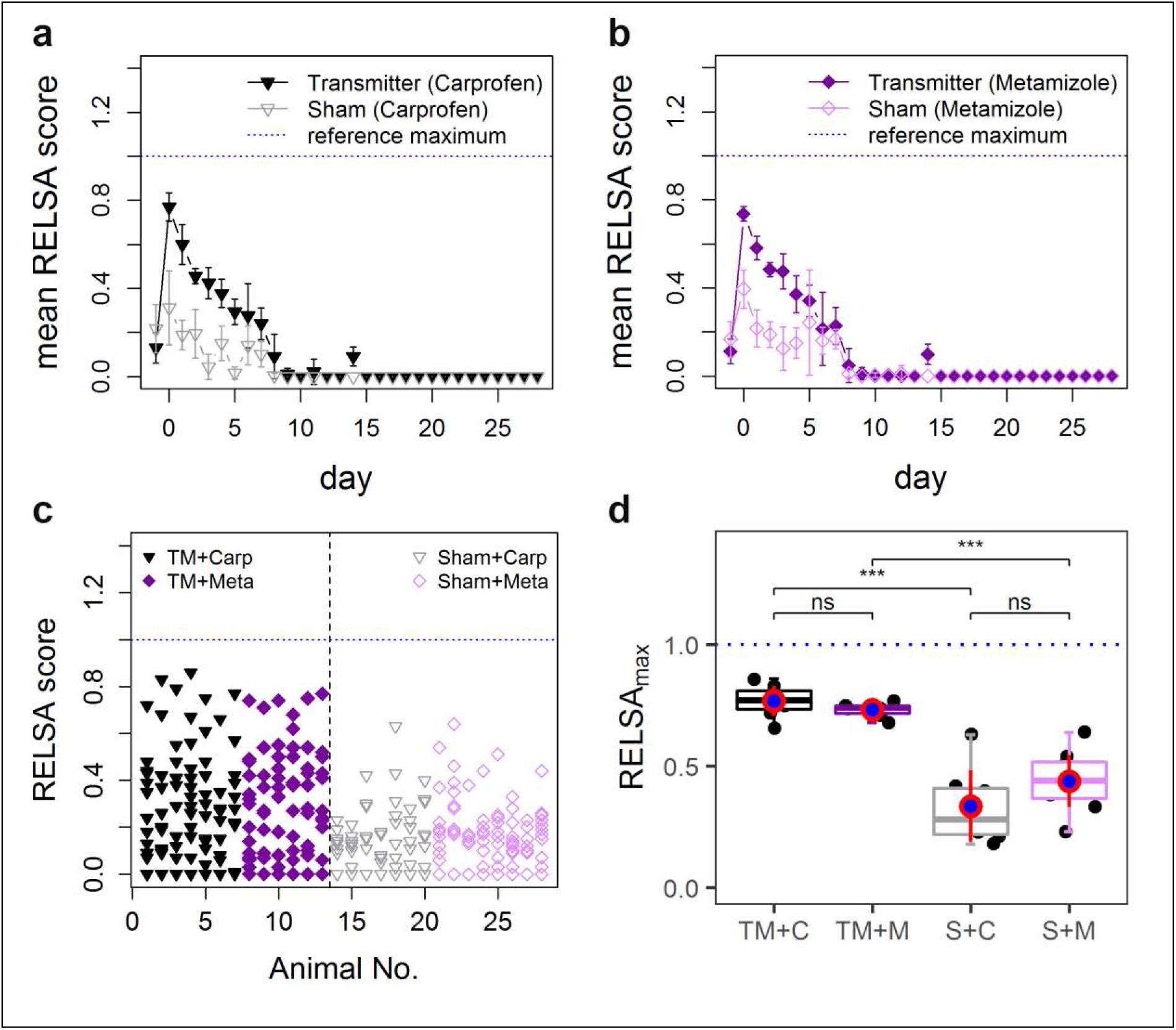
RELSA scores after transmitter implantation and sham surgery. Mean RELSA profiles for TM- and sham-operated animals treated with the analgesics (**a**) carprofen or (**b**) metamizole. In both analgesic regimen groups, the highest RELSA scores were reached on the post-op day (TM_Carp_=0.77, sham_Carp_=0.31 and TM_Meta_=0.74, sham_Meta_=0.31, respectively), followed by a decrease towards zero before postoperative day 10. (**c**) The RELSA scores of individual animals (each triangle or square represents one animal of the respective group) in both groups differ in maximum values (RELSA_max_). (**d**) Pairwise comparisons of the RELSA_max_ values in each treatment group. Comparing analgesic regimens within the TM and sham experiments, there are no significant differences. Between the transmitter and sham experiments, the differences indicate that the RELSA_max_ values are suited for experimental group comparisons (p_adj_≤0.001).

RELSA scores for single animals (Fig. 4c) highlighted the importance of individual assessments, especially because the maximal severity grades were easily detected for each individual and independent of the time course. The maximum achieved RELSA values (RELSA_max_) best reflected the overall experienced severity for the respective animals and thereby contributed to the RELSA reference model (Table 1). RELSA_max_ can also be used to delineate the overall maximum severity within a cohort of animals. Comparing RELSA_max_ on the group level revealed statistically significant differences between the two surgery groups (analysis of variance (ANOVA): p<0.0001, F_(3,24)_=25.44 with p_(TM+C|S+C)_=0.00016 and p_(TM+M|S+M)_=0.000024) but no differences between analgesia treatments (p_(TM+C|TM+M)_=0.28 and p_(S+C|S+M)_=0.20) (Fig. 4d). Additionally, there was no overlap in 95% confidence intervals between the sham and TM groups. All RELSA_max_ values in the subgroups were normally distributed in the actual data, and no significant differences within the TM groups were found (p_(TM+C|TM+M)_=0.28 and p_(S+C|S+M)_=0.20). No relevant differences in the effects of the two analgesic regimens were observed. This shows that the RELSA score, which is a representation of composite differences, allows quantification of the severity during experimental procedures in a time-dependent manner and even the determination of the maximum-experienced severity on a group level or even on a model level. This can be looked at individually or as pairwise comparisons.

### RELSA identifies the most sensitive/best performing parameters

The full set of the provided variables was used to reflect the multivariate nature of the impairment. Of course, for severity assessment in a realistic scenario, a selection of the best performing variables for a given model is desirable. We can clearly show that some single variables or combinations outperform others in this study. However, this outperformance differs from day to day. Each variable can assume a state of 0 (not chosen) or 1 (chosen). Since there are eight variables, a total of 2^8^=256 combinations were analyzed. Therefore, we tested the possible variable combinations within the pooled TM data and calculated the RELSA scores for each day. The RELSA scores were summed in a 30 by 256 matrix during the iteration steps. Finally, the individual sums were averaged using the total number of analyzed animals (n=13). The resulting RELSA performance score for each variable combination across postoperative days is shown in Fig. 5a. On the post-op day, the best performing RELSA scores in the present data were bur2h (0.96), bur2h/act (0.93), bur2h/hrv (0.9), bur2h/hrv/act (0.9) and bur2h/hr (0.89). The top 5 worst performers on the post-op day 1 were burON/temp/mgs (0.49), bwc/temp/mgs (0.48), mgs (0.29), temp/mgs (0.25) and temp (0.1) (Table 2). Using only bur2h as the best performing variable on post-op day to display individual RELSA scores revealed a similar grading of the TM vs sham groups regarding the maximum values (Fig. 5b). This underscores the rationale for selecting the best informative parameters for a given model. However, this parameter alone does not reflect the welfare state over the whole time course of this study.

**Table 2.**
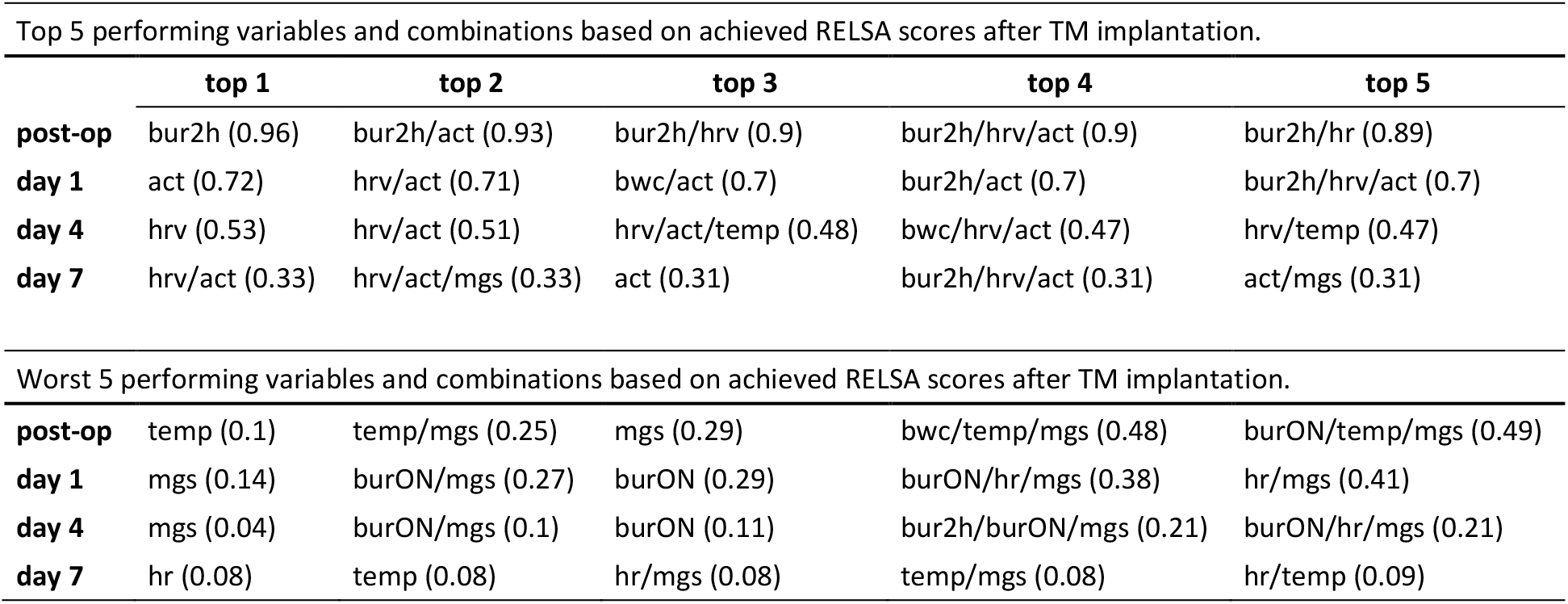
Top/worst-performing variables and combinations in the RELSA algorithm for the TM data.

**Figure 5.**
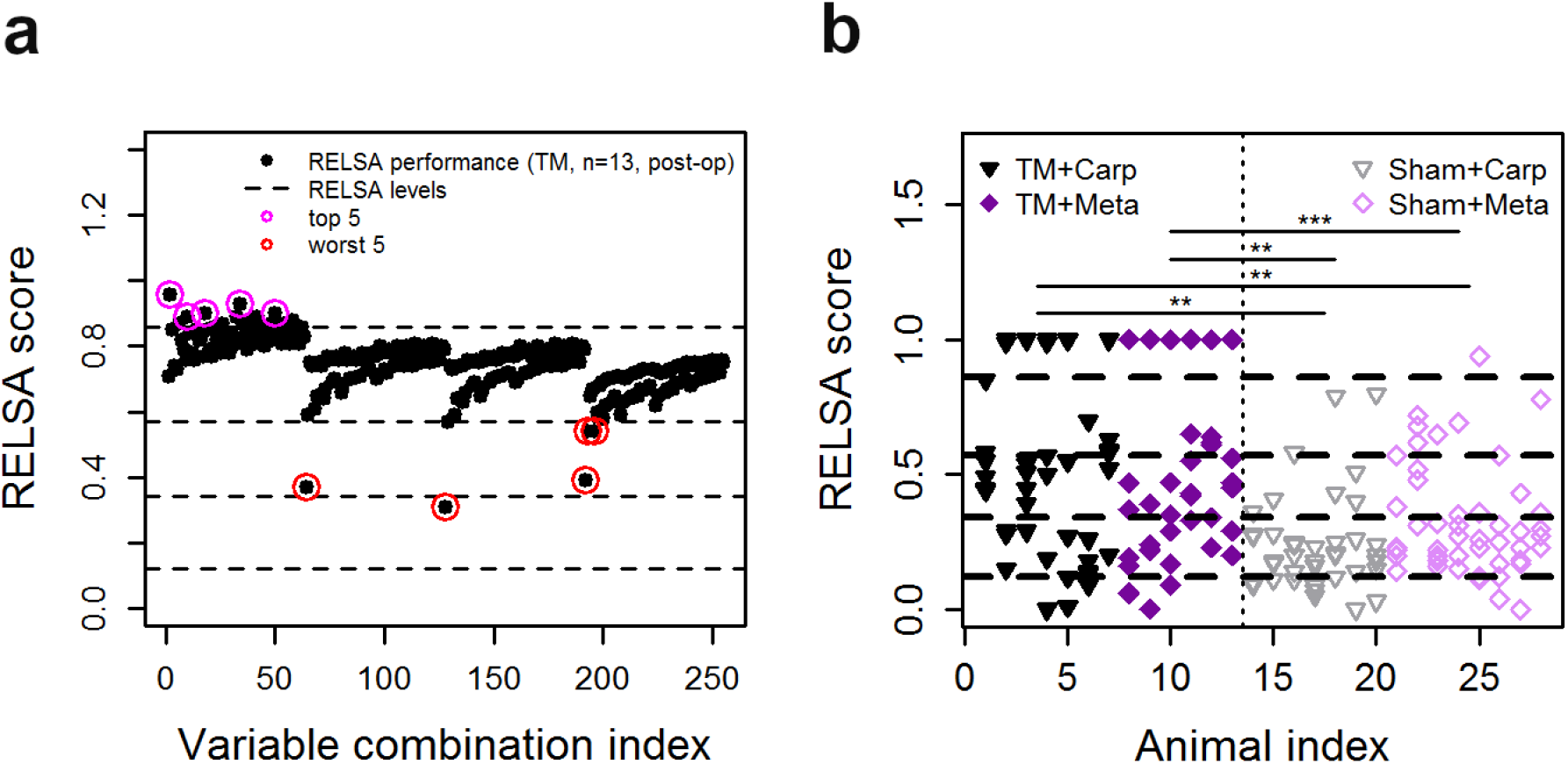
Sensitivity analysis of the average RELSA performance across postoperative days. (**a**) RELSA scores relying on combinations of the eight input variables including body weight change (bwc), Mouse Grimace Scale (mgs), burrowing behavior (bur: 2h= start before dark phase, ON= overnight), heart rate (hr), heart rate variability (hrv), body temperature (temp), and general activity (act) of transmitter (TM)-implanted mice led to 256 different combinations. High-performing variables or combinations are indicated by magenta circles, and low performing variables are highlighted red. The general performance seems to cluster locally, e.g., when good performing variables such as bur2h (0.96) are combined with low performers such as mgs (0.29) and temp (0.1). There are only a few combinations consistently showing severity on the post-op day (e.g., bur2h/hrv/act (0.9)) with high values (>0.9). Any other combination results in lower average RELSA values and is, therefore, less sensitive. (**b**) Individual RELSA profiles using only the bur2h variable. The TM group consistently shows high RELSA scores in the range [0.85; 1], whereas in the sham group, a larger range is seen [0.36; 0.94]. Within the TM- and sham-operated groups, no significant differences were detected, whereas significant differences were detected between the TM- and sham-operated subgroups.

### The clustering of RELSA_max_ scores reveals objective severity levels

In addition to the data for building the RELSA reference set from TM-implanted mice, we evaluated RELSA performance as a tool for severity comparisons between models by including data from three additional animal studies (colitis, stress, sepsis). All included studies recorded data for the following five variables: heart rate, heart rate variability, temperature, activity, and body weight. Each study was analyzed using the RELSA methodology and was therefore referenced against the data set from the TM-implanted mice. This way, the overall context allowed the comparison of studies in terms of general model severity. For this, we used the individual RELSA_max_ values as previously described to assess the maximum achieved severity for each animal in these studies. With these data, we used a *k*-means cluster analysis to segment the ordered univariate RELSA_max_ outputs into distinct clusters. We estimated the number of clusters heuristically to *k*=4 using scree analysis (Fig. 6a, b). The resulting borders of the clusters are shown as dashed lines in Fig. 6c and d. The four RELSA_max_ cluster levels were L1<0.27, L2<0.59 L3<0.82, and L4<3.49.

**Figure 6.**
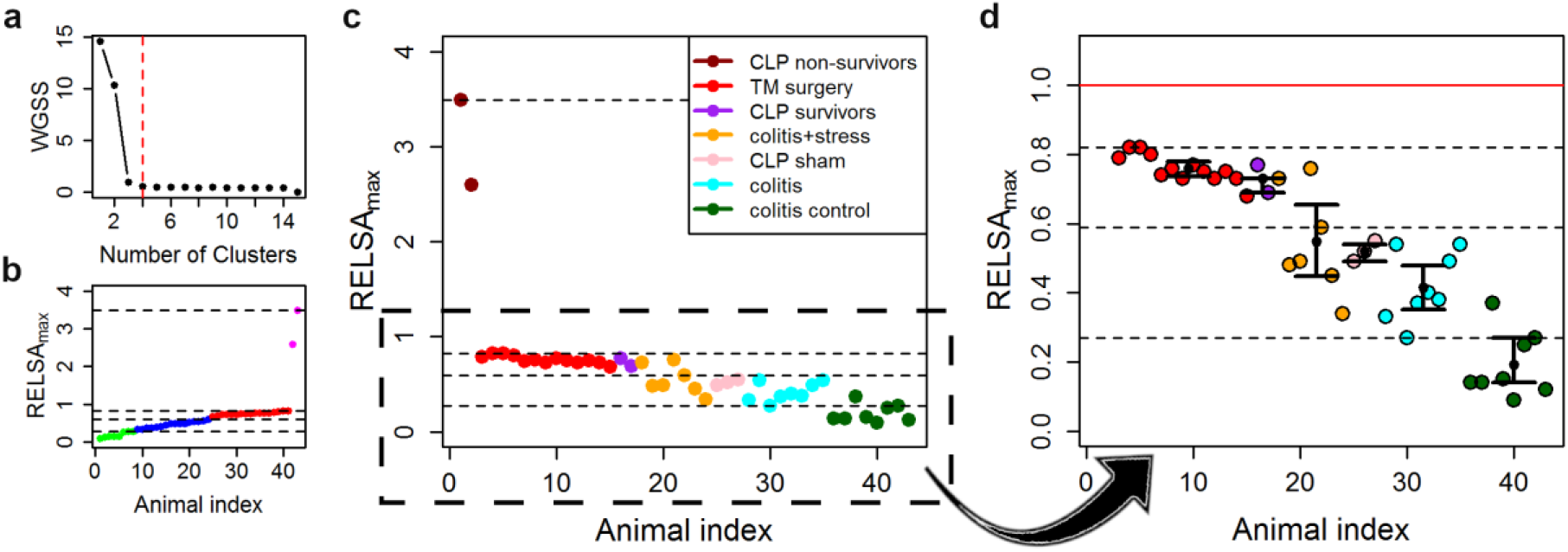
RELSA cluster analysis and severity categorization of distinct mouse models. For comparisons of the mouse models for surgery (TM implantation), sepsis and colitis/restraint stress, the variables body weight change (bwc), heart rate (hr), heart rate variability (hrv), body temperature (temp) and general activity (act) were used. (**a**) Scree plot analysis of the number of clusters against the within-groups sums of square (WGSS) values resulted in a number of clusters for the *k*-means analysis of *k*=4 (red dashed line). (**b**) *k*-means clustering of the RELSA_max_ values from all study subgroups revealed 4 cluster borders for the classification of severity grades. Clusters were found at RELSA_max_ levels: L1<0.27, L2<0.59 L3<0.82, L4<3.49. (**c**) Projection of the individual RELSA_max_ values into the cluster space. Study subgroups are color-coded. The red-colored points for the TM surgery represent the RELSA reference data and define the reference level for the algorithm. Two animals from among the CLP nonsurvivors exceeded this reference level, reaching large values (RELSA_max_ ≥2.5). (**d**) The bootstrapped cluster centers with 95% confidence intervals show that except for the data from the colitis + stress model, the intervals remain within the identified cluster levels. Individual animals in the colitis control group showed increased severity due to a drop in activity.

We used data from mice suffering from colitis induced by dextran sulfate sodium (DSS), colitis + stress where the animals received DSS and were additionally subjected to immobilization stress on 10 consecutive days for 1 h per day, and corresponding colitis control animals treated only with water. Furthermore, we used data from mice submitted to cecal ligation puncture (CLP) surgery for sepsis induction and the corresponding sham-operated animals (CLP sham). Here, the data were divided into CLP survivors and nonsurvivors. The above-described cluster levels enabled a ranking of the respective animal models regarding the severity that was experienced. Cluster analysis revealed the highest severity level for CLP nonsurvivors, followed by a cluster of TM-implanted animals (which were the RELSA reference set) and CLP survivors. Lower severity clusters were formed by data from animals suffering from colitis and stress, colitis alone and CLP sham-operated animals. Data from colitis control animals were allocated to the lowest RELSA cluster (Fig. 6c, d). Furthermore, we investigated how stable the RELSA_max_ distributions were in terms of their mean values and cluster positions. Some studies or subgroups involved small sample sizes. Therefore, we applied 10000-fold bootstrapping to assess the 95% confidence intervals of the RELSA_max_ centroids. Except for the colitis + stress study, the confidence intervals remained within their relative *k*-means cluster levels. The confidence interval for the colitis control group did not overlap with any other higher-level confidence interval.

### Model-specific parameter contributions to the general RELSA severity estimation

RELSA curves from the individual animals over time displayed the generalized biological variation that occurs during severity monitoring. Individual animals deviated from the group mean (Fig. 7a, b). This enabled individual severity monitoring. We used radar charts to quantify the contribution of single parameters to the RELSA_max_ scores (Fig. 7c). For data from the TM-implanted animals (Fig. 7b), it became obvious that immediately after surgery, all variables except for temperature contributed to the overall detected severity. Over time, some parameters returned to their baseline positions, but hrv and act remained contributors to an elevated RELSA score. In the case of the CLP model, the RELSA score was dominated by the large differences in the temperature variable. However, the other parameters, except for body weight, contributed to the overall RELSA score (Fig. S3). For the CLP study, the time variable is hours and not days (Fig. S4a, b). Therefore, the body weight variable was not flexible enough to indicate rapid impairment of the animals in a manner similar to, e.g., the temperature. Interestingly, in CLP sham animals (Fig. S4c), activity was the most active variable, but temperature and heart rate also contributed to the RELSA score. In animals suffering from colitis with (Fig. S4d) and without stress (Fig. S4e), activity was the dominating variable over the first days, but on day 7, body weight became more relevant. As expected, radar charts with data from colitis control mice showed no relevant changes within any of the observed variables (Fig. S4f).

**Figure 7.**
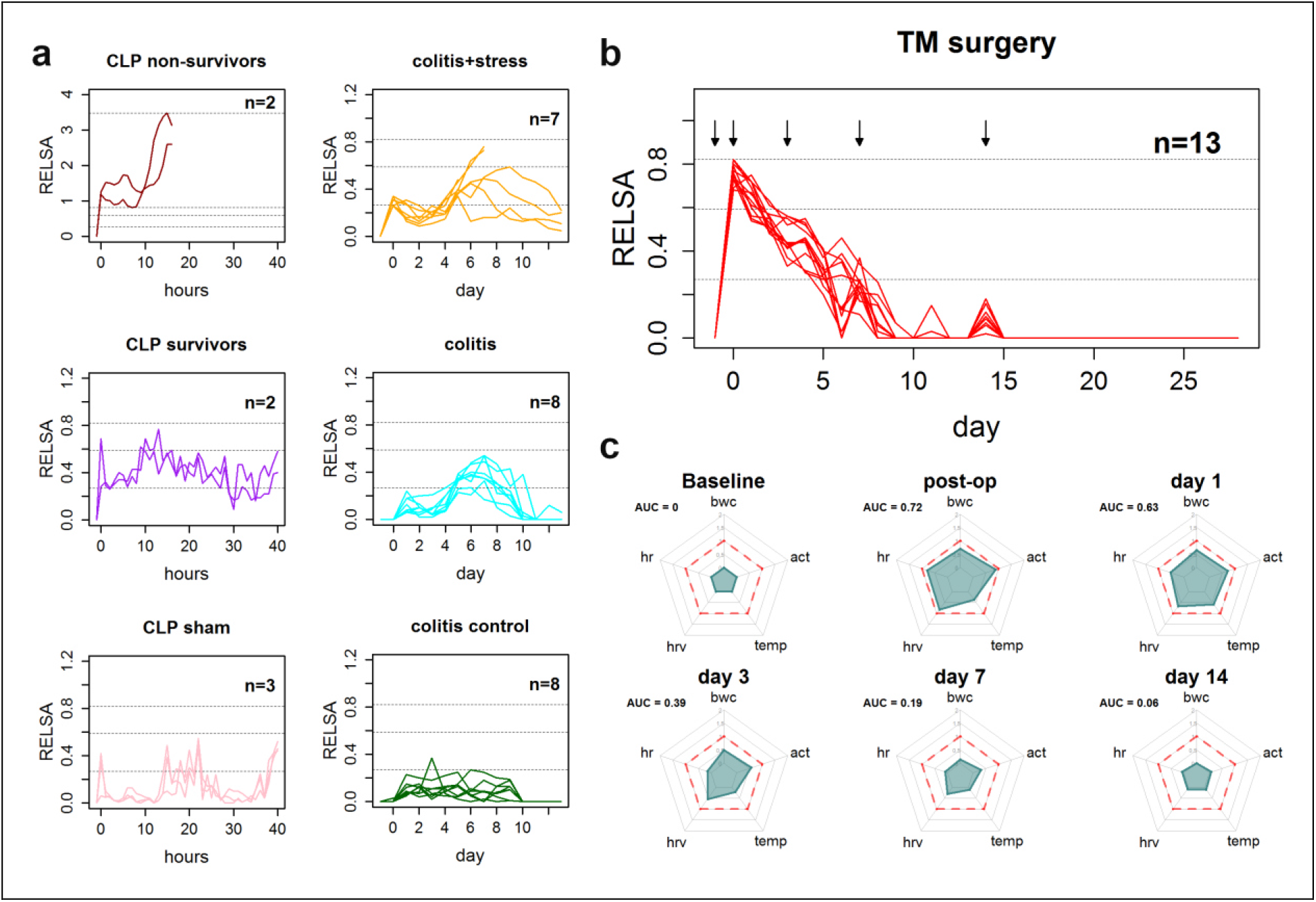
Individual RELSA time courses and variable development over the course of recovery. (**a**) Time course of RELSA scores for the animals from the cecal ligation puncture (CLP) study (left side) including the CLP nonsurvivors (brown lines), CLP survivors (purple lines) and CLP sham-operated animals (pink lines). On the right side, RELSA score development for the mice from the colitis+stress study (yellow lines). Here, the broken lines represent two euthanized animals due to meeting the endpoint criterion of 20% body weight loss. Turquoise lines represent mice suffering from colitis without additional stress treatment, and RELSA scores from the corresponding control animals are shown in green lines. (**b**) RELSA flow for the animals in the TM surgery reference model. Although the individual variance in experienced severity is high, the general direction of severity development is visible. Some animals also recover faster than others. The black arrows indicate the days for which radar charts in (**d**) for the individual variable (body weight change (bwc), Mouse Grimace Scale (mgs), burrowing behavior (bur: 2h= start before dark phase, ON= overnight)) contributions, as mean RELSA weights (R_w_), are shown. The dashed red line in the radar charts (d) indicates the RELSA reference level of 1.

## Discussion

Evidence-based severity assessment is increasingly becoming indispensable in animal research. From a researcher’s point of view, it enables the best possible monitoring of the welfare state. From an ethical point of view, it is the prerequisite for a refinement of experimental procedures leading to a minimal burden for animals and, in unity, provides a basis for high-quality data. From a legal point of view, ensuring animal welfare and severity assessment is mandatory in many countries, e.g., in all EU member states^4^. The large number and diversity of animal models and the lack of validated methods hinder clear definitions of severity categories^16^. This has multiple consequences, ranging from legal uncertainties for scientists and authorities to a potential bias in rating the prospective severity of the animals in their studies.

We have developed a tool that enables evidence-based severity assessment. With the algorithm presented, an arbitrary number of outcome variables can be used to compute a composite score for welfare assessment and severity grading^17–19^. To our knowledge, this is the first attempt in preclinical science to combine phenotypical data using matrices of standardized differences to weigh variable contributions as a means for obtaining a measure for relative severity grades. This contrasts with current standards using human judgment to generate numerical scores for assessing welfare. Using the approach presented, we have also shown that variables differ in performance and sensitivity and, therefore, strengthen the concept of a multimodal severity assessment. Finally, the RELSA algorithm enabled the quantitative comparison of distinct animal models with regard to severity levels, which leads to the speculation that it will do so in human patients as well.

### Composite scoring

When developing RELSA, we aimed at a quantitative grading of severity while methods at hand are characterized by qualitative scoring. The principle of composite scoring is based on systems utilized for clinical monitoring and risk assessment in human medicine. One example is the Acute Physiology And Chronic Health Evaluation (APACHE II) score, which was first reported in 1985. The APACHE II score comprises 12 physiological and laboratory parameters with an additional weighting for age and preadmission health status to predict the risk of death^20,21^. In contrast, the Sequential Organ Failure Assessment (SOFA) score, which was established in 1996, consists of 6 different scores assessing distinct organ dysfunction and failure^22,23^. The score describes the status of morbidity and critical illness but does not predict the outcome. Currently, the SOFA score is being used in the severity assessment of COVID-19 patients to characterize mortality among intensive care unit (ICU) patients^24^.

In veterinary medicine and laboratory animal science, there are various composite scores available, e.g., the clinical severity index for acute pancreatitis in canines^25^, composite behavior scores for pain assessment in rodents^26,27^, or composite measure schemes for rat epilepsy models^28^. These elaborated systems are generally tailored to model-specific characteristics and defined facets of severity, hindering their broad applicability across scientific fields. This results in a lack of comparability based on objective criteria. Additionally, modeling severity based on mortality outcomes –in line with increasing refinement measures over the last decades– is challenging in animal research, as euthanasia upon meeting the earliest possible humane endpoint criteria outweighs the study of other predictors of death. In a study evaluating the sickness behavior score (SBS) in a rat CLP model, the authors reported mortality in CLP nonsurvivors, which was associated with elevated levels of SBS, whereas the SBS in surviving animals returned to a mild level^29^. Similar results were observed in the present study, showing CLP nonsurviving mice in the highest severity levels and survivors in the moderate severity levels. In contrast to the SBS, RELSA severity was computed by a *k*-means algorithm to detect clusters in data coming from multiple models.

### Parameters

To create a more generalized severity assessment score, we used a system that can potentially combine any measurement or variable from the clinical and behavioral examination. This takes into account the multidimensional nature of severity, reflecting not only pain and distress but also affective emotional states. Therefore, the chosen parameters for severity assessment should be multimodal^30^. This concept is supported by growing evidence in the literature. In a study assessing severity during a chronic pancreatitis model, it was shown that the combination of multiple variables improved the sensitivity of read-out parameters ^14^. In the present study, we used a comprehensive panel of methods to monitor the welfare of animals after various experimental procedures, with TM implantation as a use case. To exclude selection bias, we calculated the models’ severity level with a full set of available variables: body weight change, burrowing behavior, MGS score and telemetry-derived parameters including hr, hrv and temperature. These parameters were selected based on increasing evidence of their suitability in various model systems as well as several round-table discussions within our German Research Foundation (DFG)-funded research consortium 2591, which focuses on severity assessment in animal-based research (www.severity-assessment.de)^7,8,31^.

We observed that even though some variables showed high sensitivity towards the implantation procedure, they only showed strong changes over a short time frame. The most prominent example here is the bur2h variable. Burrowing is a highly motivated behavior of mice and is known to be impaired under painful conditions or in mouse models of anxiety and schizophrenia^32,33^. In this study, burrowing was highly sensitive in detecting changes in welfare but only immediately after TM implantation. Likewise, bwc sensitively indicated the impact of TM surgery but quickly recovered within 4 to 6 days after the operation. Body weight is considered one of the most critical parameters in classic clinical scoring in rodents^12^. However, monitoring body weight as a severity assessment parameter was shown to be model-specific and should be used in combination with other parameters^12^. Similarly, using mgs, only short-term effects were detected within 180 minutes after implantation (not shown). On a daily scale, the mgs variable played no role in indicating severity.

In contrast, the telemetry-derived parameters hr and hrv showed strong changes on the post-op day but also indicated a longer-lasting impact on animals, suggesting an extended recovery period (up to day 14). Telemetry is a frequently used method in biomedical research. It has been shown that hr and hrv are parameters indicating distress and pain^34,35^, and hr and body temp serve as critical parameters in sepsis studies^36^.

This leads to the assumption that the various parameters reflected different facets of severity (e.g., pain) better than others or that the animals did not experience the particular facets after a while. However, this question remains elusive, and the results of the present study underscore the need for a combination of parameters, including physiological parameters, to fully assess the severity situation.

This time-dependent contribution of parameters was quantitatively demonstrated by the sensitivity analysis, in which combinations of the variables were used to find the best performing set on each day of the analysis. This analysis clearly revealed that there was no general set of variables depicting severity over the whole monitoring period. This was suggested by PCA variable contributions; however, PCA generalizes over all variables and input times. This process does not include the specific analytical qualities of each variable on every analyzed day. Depending on the overall loss in variance representation (percentage of information contribution per principal component), the actual severity prediction depending only on single contributing variables is poor. It appears that different variables are rather useful for different analytical purposes (e.g., acute pain vs. long-term impairment).

### Using RELSA for comparisons

The RELSA results did not necessarily follow a normal distribution. In the TM model, the outcomes were skewed towards the time point with the dominant deviations (post-op day). Since the exact maximum depends on the animal model under observation, better choices for comparisons are the individual RELSA_max_ values. The most extreme values reveal more about the maximally achieved severity than the mean. If the animal model is stable and the results do not suffer from too much variance, the resulting RELSA_max_ values can be used, e.g., in animal model comparisons (Figs. 3 and 4).

Comparing the RELSA_max_ values revealed that TM implantation exhibited higher severity than sham operations. However, sham operation also shows some level of severity. The small peak in the RELSA score on day 14 after TM implantation (Fig. 3 a, b) demonstrates how sensitive the algorithm is towards value changes. Here, the RELSA score was not zero like the rest of the variables, but it was slightly elevated due to some minor variation in the hr variable (RELSA_hr,14_=0.021 (SD 0.06)). If there are changes in the measured values, the RELSA score will adequately reflect this. An overall effect of the chosen analgesics was not observed, leaving the search for an ideal treatment for future studies.

### RELSA algorithm validation

To validate the RELSA algorithm, we used data from models with different forms and grades of impairments. An acute DSS-colitis model, an acute DSS colitis in combination with repeated restraint stress model and a CLP sepsis model were assessed. Fig. 4c shows that the RELSA_max_ scores remained within the moderate frame of the 4 *k*-means cluster levels and did not exceed the RELSA level of 1, with the exception of the CLP nonsurvivors. The colitis RELSA_max_ values reliably clustered in level 2, indicating a lower severity for the DSS-colitis model compared to the TM-implantation study. However, in the colitis study, 9 animals had to be euthanized because the humane endpoint (max. of 20% weight loss) was reached. This had been set to ensure that animals experience a maximum of moderate severity levels according to the project authorization. Although the RELSA values indicated increased suffering, they also imply that the animals may have been euthanized too early, challenging the use of a 20% loss of body weight as an objective endpoint to ensure moderate severity levels. Even though the humane endpoint for a single variable was reached, the remaining variables did not support a general increase in overall suffering in relation to the reference set.

Data from the CLP study revealed very high RELSA scores for the animals that did not survive the procedure (RELSA_max_ ≥ 2.60) and lower values for the surviving and sham animals (RELSA_max_ < 1). The main factor responsible for the high scores was a large decrease in temperature, but hrv and act also indicated increases in severity. Here, more than one variable is pointing towards increased suffering and therefore to an increased impairment in well-being.

### RELSA principle and critical issues

RELSA enables scientists to quantify severity. In addition, it can be used to classify animals and models in qualitative frameworks, e.g., mild, moderate, and severe. For qualitative grading, data from a predefined reference set are needed, and subsequently, the severity context can be extrapolated. Trivially, the extrema (min/max) of each variable serve as ranges for the given severity context. One caveat is that researchers must provide some sort of estimation about the quality of severity for the reference set, a step that involves human judgment. However, once defined, a new experiment can be used for severity quantification with regard to the reference set. This concept is new to the field and allows an evidence-based comparison of models within actual statutory provisions and guidelines.

In addition to providing context, the reference set has another purpose: it regularizes the possible ranges of the input variables. This can prove essential, as variables behave differently when animals are negatively affected. For example, a loss of 17% in body weight is generally recognized as a threat to animal health^12^. At the same time, burrowing behavior may drop to zero. In this case, a difference of 17% in one variable is equivalent to 100% in the other variable. For an optimal representation of this bias, we calculated individual RELSA weights (R_w_) as effect sizes for each variable and day, which were then used in the final score calculation.

Individual variables contribute to the final RELSA score as RELSA weights (R_w_). These weights can be considered a special form of effect size that is somewhat related to Glass’ Ҕ^37^. For the R_w_ values, however, the differences are that these values are not standardized to the standard deviation in the control group but rather to the difference of the respective variable to its maximum deviation in the reference set. This approach allows an estimation of within-animal effect sizes and measurements of a particular variable’s importance.

For the generalization of the weights in a final score, we concluded that variables with larger deviations should have more impact, while smaller deviations mostly represent noise and effects that are less prominent within a cohort. In statistics, this is followed by the root mean square (RMS) concept, e.g., in error and regression analysis. In contrast to a pure sum score, the RMS has the advantage that it directly translates to the scale of the individual weights and is considered to be more accurate in showing the best fit.

Another important issue is the sampling and measurement frequency. Body weight is detected (e.g., once per day in the morning) and burrowing behavior after a certain time (e.g., after 2 h or overnight). The sampling rates in these cases are a) not equal and b) not frequent enough to catch minute-by-minute changes. Transient changes in some variables thus appear as “all-or-nothing” parameters. They change much faster than the sampling rates so that the exact development over time cannot be seen. Although the sampling rate cannot be corrected with RELSA, the skewness in distribution can be adjusted to a certain degree by including extreme values of a reference model with known severity into the calculation. To be comparable, the RELSA algorithm requires the same reporting frame (e.g., day) in all input variables even though this can mean that the integration times are different (e.g., bur2h).

### Outlook and conclusion

RELSA was designed to assess the multidimensional severity an animal experiences under impaired welfare conditions using multivariate data. The combination of objective variables into a composite score has the advantage of unbiased severity assessment without the need for interpretation or analysis. We have shown that such a composite model can be built, tested and validated. In the future, a comparison of more animal models will lead to a severity map that can then be used to obtain a better understanding of the multivariate severity context. It will not only become much clearer to assess severity but also enable the ranking of animal models with regard to their impairment of welfare. Finally, this may also reveal more generalized or specific variables for monitoring severity. With the development of home cage monitoring systems, RELSA enables an automatic continuous assessment of the animals and thereby an early warning system helping to identify animals at risk.

## Methods

### Ethical Statement

Experiments involving surgery, DSS colitis and stress were approved by the Local Institutional Animal Care and Research Advisory Committee and permitted by the Lower Saxony State Office for Consumer Protection and Food Safety (LAVES, license 15/1905). The application for the animal experiments involving sepsis (authorization no. V54 – 19 c 20/15 - F152/1016) was approved by the local Ethics Committee for Animal Research (Darmstadt, Hesse, Germany). All procedures were carried out in accordance with the German law for animal protection and the European Directive 2010/63/EU.

### Animals, housing conditions and husbandry

Female C57BL/6J mice undergoing surgery only (transmitter implantation and sham groups), DSS colitis or stress induction were obtained from the Central Animal Facility, Hannover Medical School, Hannover, Germany. For the sepsis study, male C57BL/6N mice were obtained from Charles River Laboratories, Sulzfeld, Germany (for an overview on the studies, mice and animal numbers see Table S6). The mice were free of the viral, bacterial and parasitic pathogens listed in the recommendations of the Federation of European Laboratory Animal Science Association^38^. Their health status was monitored by a sentinel program throughout the experiments. The mice were housed at the Central Animal Facilities of the MHH (surgery, colitis, stress groups) in macrolon type-II cages (360 cm^2^; Tecniplast, Italy), which were changed once per week. Cages were bedded with autoclaved softwood shavings (poplar wood; AB 368P, AsBe-wood GmbH, Buxtehude, Germany), paper nesting material (AsBe-wood GmbH, Buxtehude, Germany) and two cotton nesting pads (AsBe-wood GmbH, Buxtehude, Germany). Room conditions were standardized (22 +/− 1°C; humidity: 50%-60%; 14:10 h light/dark cycle). Mice were fed standard rodent food (Altromin 1324, Altromin, Lage, Germany) *ad libitum,* and autoclaved (135°C/60 minutes) distilled water was provided *ad libitum*. The mice were handled by two female persons. For the sepsis experiments, the mice were housed at the animal facility of Fraunhofer IME-TMP, Frankfurt, Germany in IVC cages (501 cm^2^; GM500, Tecniplast, Italy), which were changed once per week (but never during the course of sepsis). These cages were bedded with softwood shavings (H0234-200, ssniff Spezialdiäten GmbH, Germany), paper nesting material (Sizzlenest, H4201-11, ssniff Spezialdiäten GmbH, Germany) and a mouse igloo (#13100 Plexx BV, Netherlands). Room conditions were standardized (22 +/− 2°C; humidity: 45%-65%; 12:12 h light/dark cycle including a 30-min twilight phase at the beginning and end of the light/dark phases). The mice were fed standard rodent food (V1534-000, ssniff Spezialdiäten GmbH, Germany) ad libitum, and tap water was provided ad libitum.

All mice were randomly allocated to the testing groups and habituated to the experimental environment before the surgical procedure.

### Data acquisition and experimental setup

#### Transmitter implantation

The mice for the surgery, colitis and stress studies were aged 9-10 weeks. Transmitters (ETA-F10 or HD-X11; DSI, St Paul, MN, USA) were aseptically implanted into the intraperitoneal cavity with electrodes placed subcutaneously for a bipolar lead II configuration under general isoflurane anesthesia. General anesthesia was induced in an induction chamber (15 × 10 × 10 cm) with 5 vol% isoflurane (Isofluran CP^®^, CP Pharma, Burgdorf, Germany) and an oxygen flow (100% oxygen) of 6 l/min. After confirmation of the absence of the righting reflex and removal from the chamber, anesthesia was maintained via an inhalation mask with 1.5-2.5 vol% isoflurane and an oxygen flow of 1 l/min. The depth of anesthesia was monitored and controlled by means of the corneal and eyelid reflex. To protect the eyes from drying out, the eyes were moistened with eye ointment (Bepanthen®, Bayer AG, Leverkusen, Germany). After reaching full anesthesia, the surgical area was shaved, and the mice were placed in the surgical field in dorsal recumbency with the head towards the surgeon. During the entire duration of the anesthesia, the mice were placed on a heating pad at 37.0 ± 1.0°C to prevent hypothermia. For local anesthesia at the incision sites, xylocaine (Xylocain^®^ Pumpspray Dental, AstraZeneca GmbH, Wedel, Germany) was used. The mice that underwent only surgery received either preoperative 200 mg/kg metamizole subcutaneously (s.c.) and postoperative 200 mg/kg metamizole orally via the drinking water until day 3 or preoperative 5 mg/kg carprofen s.c. and postoperative 2.5 mg/kg s.c. every 12 h until day 3. The mice that underwent additional colitis or stress induction were treated using the metamizole analgesia regimen.

In the CLP study, mice aged 12 to 14 weeks were anesthetized via s.c. injection of 120 mg/kg in 10 ml/kg ketamine (Ketaset^®^, Zoetis Deutschland GmbH, Berlin, Germany) and 8 mg/kg in 10 ml/kg xylazine (Rompun^®^, Bayer Vital GmbH, Leverkusen, Germany). Perioperative management was the same as described above. The blood pressure catheter was placed in the left carotid artery and positioned so that the gel-filled sensing region of the catheter was approximately 2 mm in the aortic arch. The telemetry transmitter device body was placed along the lateral flank between the forelimb and hindlimb, close to the back midline. Biopotential ECG leads were tunneled subcutaneously to achieve positioning analogous to lead II in human ECG. For postsurgical analgesia, 200 mg/kg metamizole s.c. (Novaminsulfon 1000 mg Lichtenstein, Zentiva Pharma GmbH, Frankfurt/Main, Germany) was administered at the first signs of waking up. For postsurgical analgesia, 5 mg/kg carprofen (Rimadyl, Zoetis Deutschland GmbH, Berlin, Germany) was administered s.c. on the evening of the day of the surgery and on the morning and evening of day 1 and day 2 after surgery. After fully recovering from the anesthesia, mice were put back into their home cage, and the continuous data acquisition of all physiological parameters began immediately. Mice were randomly allocated to the testing groups and habituated to the experimental environment before the surgical CLP or CLP sham procedure.

### Sham surgery

Sham-operated mice were used as controls for assessing the severity of transmitter implantation and underwent an aseptic laparotomy without transmitter implantation (sham mice) under the same conditions as described above (surgery studies), including anesthetic and analgesic regimens.

### Burrowing behavior

One week before intraperitoneal transmitter implantation or the corresponding sham surgery, the mice were housed pairwise in type ll macrolon cages filled with aspen bedding material (AsBewood GmbH, Buxtehude, Germany) and two compressed cotton nesting pads (AsBewood GmbH, Buxtehude, Germany). On days five and four before surgery, the burrowing apparatus was provided to the animals to train burrowing behavior^39^. Baseline measurements were taken on days two and one before surgery. A 250-ml plastic bottle with a length of 15 cm, a diameter of 5.5 cm and a port diameter of 4 cm was used as a burrowing apparatus. It was filled with 140 g +/− 1.5 g of the standard diet pellets of the mice (Altromin1324, Lage, Germany).

For burrowing testing after surgeries (1^st^, 2^nd^, 3^rd^, 5^th^ and 7^th^ night after surgery), mice were single housed in a type-II macrolon cage with autoclaved hardwood shavings. The burrowing bottles were placed in the left corner. In the right corner, half of the used nesting material from the home cage was provided as a shelter. The tests started three hours before the dark phase, and after two hours, the content of the burrowing bottles was weighed (bur2h). The bottles containing the remaining pellets were placed back into the cages and weighed again the next morning (burON).

### Mouse Grimace Scale

For mouse grimace scoring, the mice were individually placed in 9 × 5 × 5 cm cubical boxes. The two long sides of the boxes consisted of red plexiglass, and the other parts consisted of transparent plexiglass. The front side contained three air holes at the bottom, and the back side contained several air holes. Mice were recorded with a Canon EOS 750 D camera for 12 minutes^40,41^.

Video footage was analyzed using the VLC Media Player^®^. Eight pictures per time point and animal in their respective treatment conditions (n=4000) were extracted in regular intervals or when a face exhibition was observed. The pictures were grabbed via screenshot and cut out using Paint^®^ so that only the mouse’s face was visible. The pictures were randomized with the ‘RandomSlides’ PowerPoint Tool and evaluated by two blinded scorers. According to Langford *et al* 2010^42^, they scored five “action units” (AU) (orbital tightening, nose bulge, cheek bulge, ear position and whisker change) per picture with values of “0” (AU is not present), “1” (AU is moderately present) or “2” (AU is severely present).

Baseline MGS scores were determined from the two consecutive days before surgery. After surgery, mice were filmed for 30 and 180 minutes and on the 1^st^, 2^nd^, 3^rd^, 5^th^ and 7^th^ day at 9 am when the burrowing testing was finished. Afterward, the mice were transferred back to their group home cages.

### Cecal ligation and puncture surgery

At the earliest 6 days or after reestablishing a regular circadian rhythm after the surgical implantation of the telemetry transmitter device, male C57BL/6JN mice were used for the CLP experiments. The CLP surgery and the subsequent start of the experiments were conducted in the morning to control for circadian variations. The mice were weighed, and 30 minutes prior to surgery, 0.05 mg/kg buprenorphine was injected s.c. (Bupresol^®^ 0.3 mg/ml, CP-Pharma HmbH, Burgdorf, Germany). The mice were anesthetized using isoflurane anesthesia (2-3% Forene^®^, AbbVie Deutschland GmbH & Co. KG, Wiesbaden, Germany) and placed on their back on the heating pad, while continuously connected to the isoflurane anesthesia. The eyes were moistened with eye ointment. For local anesthesia at the incision site, xylocaine (Xylocain^®^ Pumpspray Dental, AstraZeneca GmbH, Wedel, Germany) was used. The depth of anesthesia was checked by means of the corneal and eyelid reflex. During the entire period of anesthesia, the mice were on a heating pad at 37.0 ± 1.0°C. The abdominal cavity was aseptically opened via a midline laparotomy incision of approximately 3 cm, and the cecum was exposed. Subsequently, the cecum was 2/3 ligated (Nylon Monofilament Suture 6/0, Fine Science Tools GmbH, Heidelberg, Germany) distal to the ileocecal valve, while care was taken that the intestinal continuity was maintained. The exposed cecum was punctured twice, “through-and-through”, with a 21-gauge needle. Next, sufficient pressure was applied to the cecum to extrude fecal material from each puncture site (~ 1 mm). The cecum was returned to the abdominal cavity and placed in the upper central abdomen. Following this procedure, the peritoneum was closed with three knot fissures with nonresorbable sterile suture material (Nylon Monofilament Suture 7/0, Fine Science Tools GmbH, Heidelberg, Germany), and the upper skin layer was stapled with sterile clips (Michel Suture Clips 7.5 × 1.75 mm, Fine Science Tools GmbH, Heidelberg, Germany). For the mice undergoing a sham laparotomy, the same procedure was performed without CLP. After fully recovering from the anesthesia, the mice were put back into their home cage, after which the continuous data acquisition of all physiological parameters began immediately. The mice received 0.1 mg/kg buprenorphine s.c. three hours after surgery and subsequently every 8 h for the rest of the experiment. At the end of the experiments, mice were anaesthetized deeply with isoflurane and killed by cervical dislocation. .

### Colitis induction and restraint stress

After intraperitoneal transmitter implantation and 28 days of postoperative recovery, the female C57BL6/J mice were exposed to 0% (control; receiving water only) or 1% DSS (colitis; mol wt 36000–50000; MP Biomedicals, Eschwege, Germany) in drinking water for 5 consecutive days to induce intestinal inflammation. The mice were weighed daily, and the telemetry-derived parameters hr, hrv, activity, and temperature were recorded. A third group of mice was subjected to restraint stress (colitis + stress) in addition to DSS treatment. The mice were inserted into restraint tubes on 10 consecutive days (d1-d10) for 60 minutes (from 09:00 to 10:00 am). The restraint tubes (23-mm internal diameter, 93-mm length) consisted of clear acrylic glass with ventilation holes (8 mm diameter) and a whole length spanning 7-mm–wide opening along the upper side of the tube. The ends of the tube were sealed on one side by a piece of acrylic glass with a slot for the mouse tail and on the other end by a solid plastic ring that screwed into place. The mice were able to rotate around their axis but could not move horizontally.

### Data characterization

Before analysis, the data were brought into the tabular format required for RELSA analysis (Table S5). Eight variables were used in the calculations (body weight change (bwc), Mouse Grimace Scale (mgs), 2 h of burrowing started 3 h before dark phasebur2h), burrowing overnight (burON), heart rate (hr), heart rate variability (hrv), body temperature (temp) and activity (act)). For each variable, the data were pooled, and the effective ranges were determined (Table 1). Furthermore, Cohen’s d was calculated for each variable and each day, i.e., post-op (0), 1, 4 and 7, to compare the resulting RELSA scores with an independent measurement of effect.

### Principal component analysis (PCA)

PCA was conducted using the factoextra package in R. PCA requires complete data so that the present data were limited to the following days: baseline, post-op and postoperative days 1 to 4. For PCA calculations, all variables were scaled and centered. The principal components of the first two dimensions for all respective days were plotted, as well as the factor loadings and variable contributions.

### Relative Severity Assessment (RELSA) score calculation

The principal methodology of the RELSA calculation is depicted in Fig. 3. Quantitative input data were normalized to the range [0; 100]% with 100% as starting values (based on physiological or baseline conditions, e.g., on pre-op day (−1); Table S5).

The RELSA methodology requires a reference set. If this set has a qualitative severity attribute, the calculated scores will be in reference to that category. According to Annex XIII of the EU directive, surgical interventions under general anesthesia, such as the TM implantations or sham surgery, are categorized as “moderate” in terms of severity. Thus, the RELSA reference set quantitatively reflected this category. It uses the respective extrema of the monitored variables, thereby establishing the context within the referential severity category. For each time point (*t*), data differences from the normalized baseline for each contributing variable (*i*) were calculated. To establish the severity context, the differences were divided by the normalized maximum-reached differences in the respective variables of the reference set to yield weights (R_*w*_, *see formula 1*). For this measure, absolute differences were used. Each R_*w*_ is an expression of the similarity of an actual data point to the maximum-reached value observed in the reference set at any observed time point. This step also regularized differences in variable contributions at any given level of severity so that different scales do not skew the results. To give larger differences more weight, the final RELSA score was calculated by the root mean square (RMS) of the available R_*w*_ divided by the number of variables (N) *(see formula 2)*. Missing variables did not contribute to the RELSA score, whereas values equal or above baseline level contributed with values of zero. Furthermore, levels of severity in the reference data were calculated using a *k-*means algorithm^13^. The number of clusters was determined heuristically with a scree plot (Fig. 4a). A RELSA score of 1 means that all contributing variables for a test animal reached the same values as the largest observed deviations in the reference set with the defined level of severity (here, “moderate”).

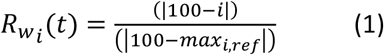

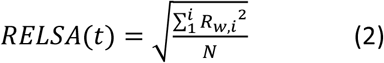

### Sensitivity analysis

For each animal, RELSA scores were calculated for every available variable and its combinations. Each variable can assume a state of 0 (not chosen) or 1 (chosen), leading to 2^n^ possible combinations. The analysis was repeated for each day to reveal the time-dependent changes for all variables and their combinations. The results indicated variable sensitivity as a measurement of average RELSA performance.

### Statistics

Data distributions were tested against the hypothesis of normality using the Shapiro-Wilk test. In the case of failed rejections, nonparametric methods were used for group comparisons (Kruskal-Wallis test) and the Mann-Whitney U-test for pairwise tests. Parametric analyses were performed using analysis of variance (ANOVA) or single *t*-tests (with Welch’s correction in case of unequal variance). For multiple comparisons, the resulting p-values were adjusted using the Bonferroni correction. The RELSA_max_ and cluster centroids were bootstrapped 10000-fold to yield mean values as well as 95% bias-corrected and accelerated (BCa) confidence intervals. With either method, the resulting p-values were considered to be significant at the following levels: 0.05 (*), 0.01 (**), 0.001 (***) and 0.0001 (****).

### Software and Packages

The algorithm was developed in R (version 3.6.0). In addition to RELSA, the following packages were used for analysis: ggplot2, factoextra, effsize, plyr, and boot. Radar charts were realized using the fsmb package. The RELSA algorithm and the raw data are available as an R package with full documentation on GitHub: https://github.com/mytalbot/relsa.

### Reporting summary

Further information on the research design is available in the Nature Research Reporting Summary linked to this article.

## Data availability

The datasets used in this study are available on LINK (will be available with publication). Further requests should be made to the corresponding author.

## Author contributions

C.H. and A.B. designed the animal study. C.H., B.S., L.W., N.W., M.H., L.K., P.J., T.K., M.C.J.H, and A.v.K. conducted the experiments, collected and annotated the data and performed the descriptive analysis. S.R.T. redesigned the data annotations, developed and coded the RELSA score, developed the R package and conducted the corresponding (statistical) analyses. S.R.T., B.S., L.W., A.B. and C.H. drafted the manuscript. All authors discussed the results and commented on the manuscript.

## Acknowledgments

This research was supported by the DFG research group FOR 2591 (BL953/10-1, BL953/11-1) and by the research funding program *Landes-Offensive zur Entwicklung wissenschaftlich-ökonomischer Exzellenz* (LOEWE) of the State of Hesse, Germany.

## Additional information

Supplementary information is available for this paper.

