## supplemental for "One score to rule them all: severity assessment in laboratory mice"

### Supplemental Material

#### Figure S1

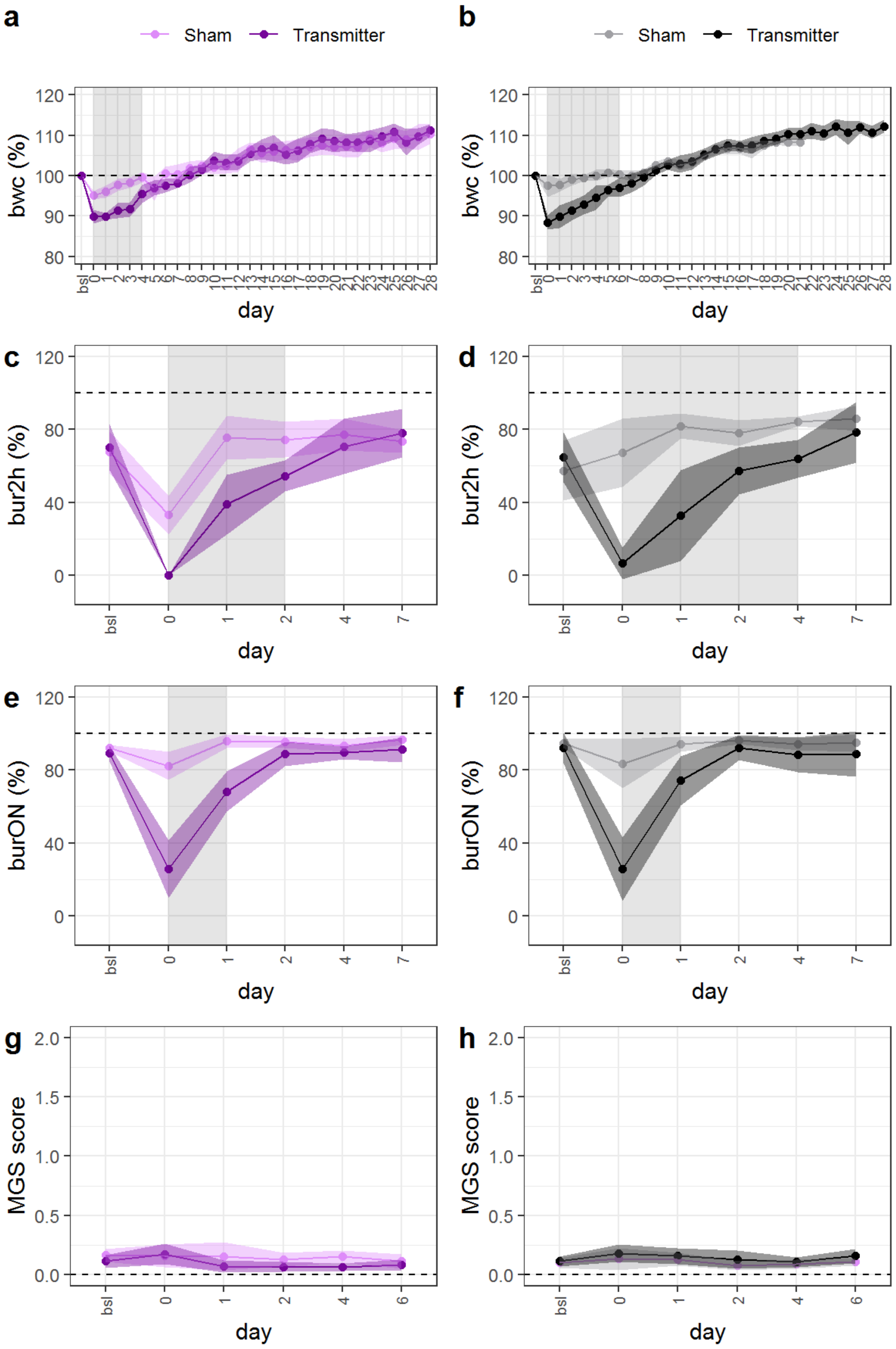


**Figure S1. Severity assessment variables after surgery.** Physiological and behavioral variables ((**a,b**) body weight change (bwc (%)), (**c,d**) burrowing after 2 h (bur2h (%)), (**e,f**) burrowing overnight (burON (%)), and (**g,h**) the Mouse Grimace Scale (MGS) score) were monitored in transmitter-implanted (purple and black lines) and sham-operated (mauve and gray) mice treated with metamizole (left) or carprofen (right) for analgesia. Brighter shades of the displayed lines represent the 95% confidence band of the respective variable. Areas with no overlap in confidence bands are marked in gray, indicating evidence for significant differences between surgical procedures.

**Figure S2**

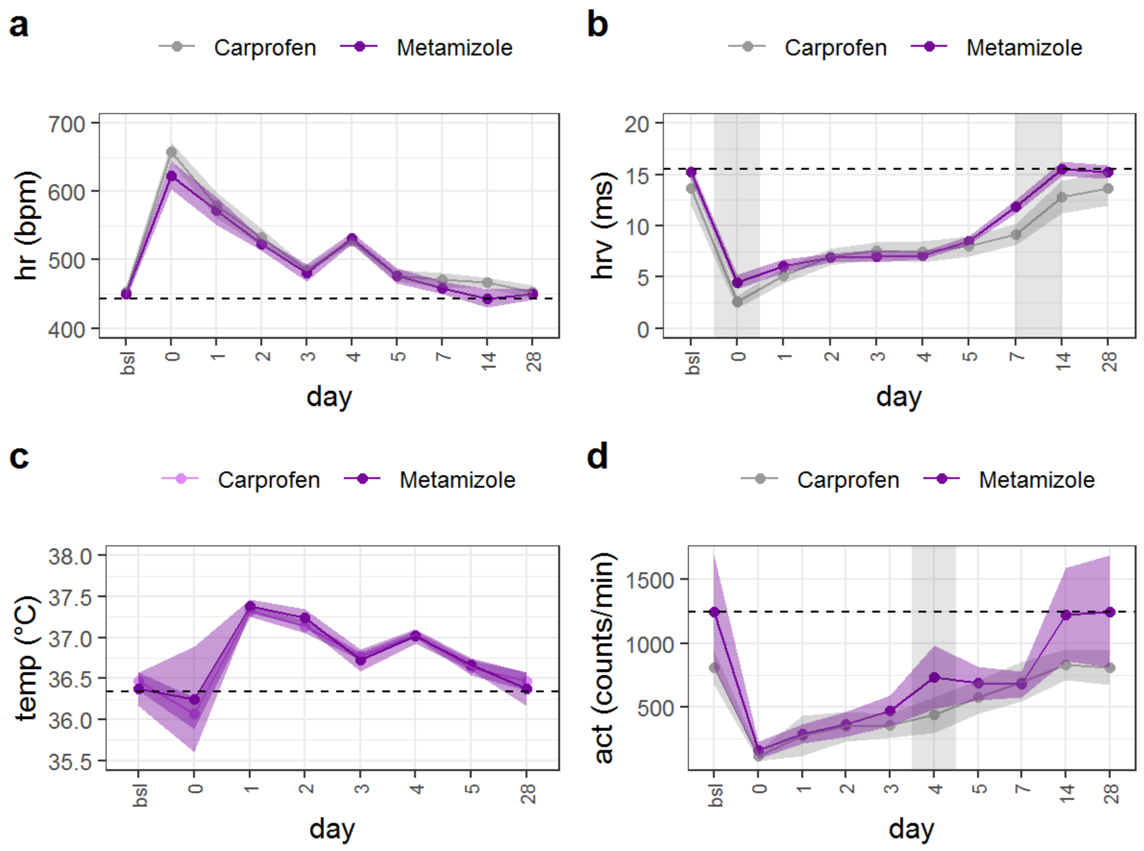


**Figure S2. Telemetry-derived variables monitored for severity assessment.** The variables heart rate (hr, in beats per minute (bpm)), heart rate variability (hrv in milliseconds (ms)), core body temperature (temp in °C) and general activity (act in counts per minute) were monitored via telemetry from transmitter-implanted mice after surgery. For analgesia, the animals were treated with carprofen (mauve) or metamizole (purple). Brighter shades of the displayed lines represent the 95% confidence band of each variable. Areas with no overlap in confidence bands are marked gray, indicating evidence for differences between the utilized analgesics.

**Figure S3**

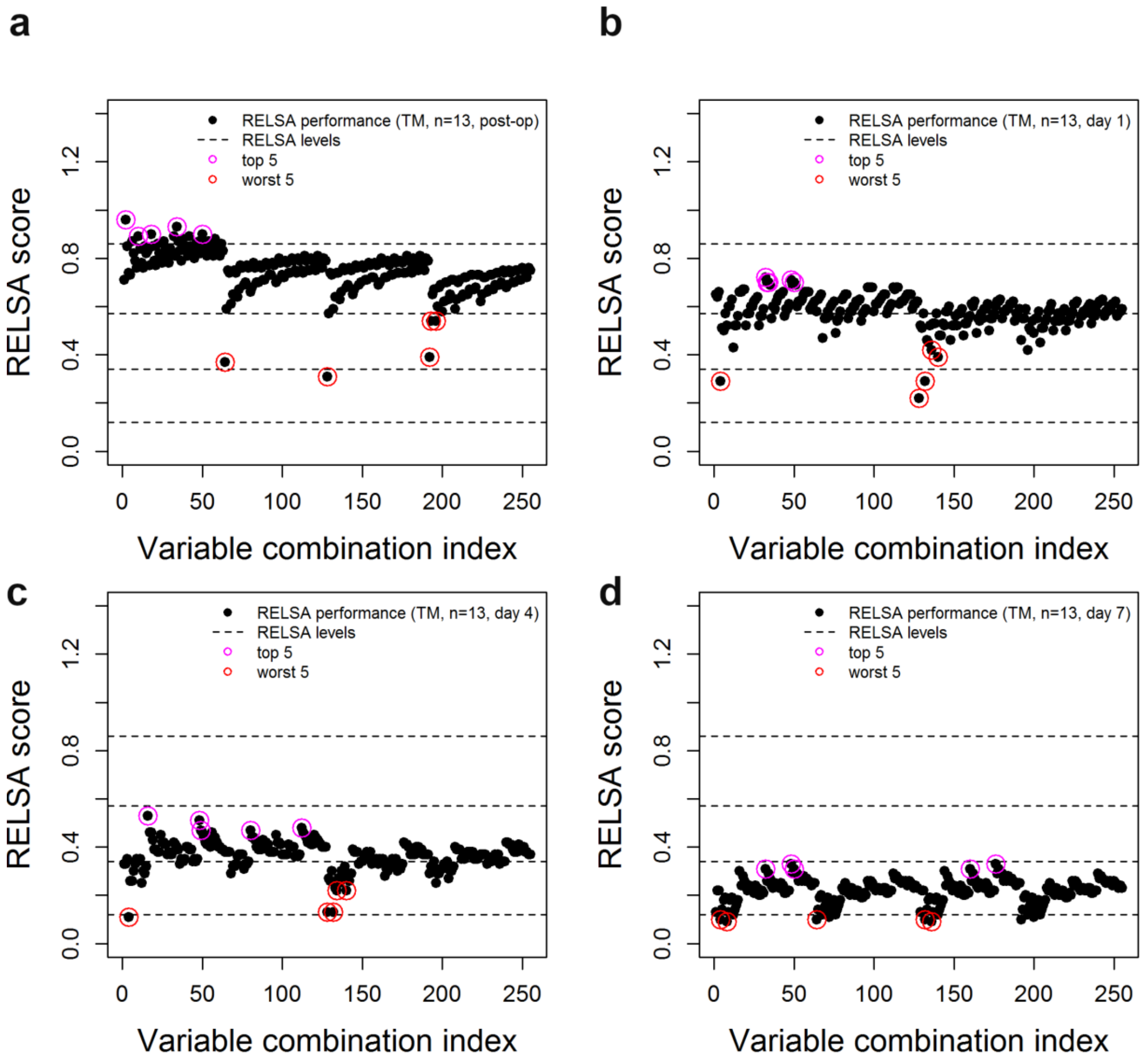


**Figure S3.** RELSA variable combinations and sensitivity analysis. For the eight input variables in the transmitter-operated animals (metamizole (n= 6) and carprofen (n= 7) treatment; pooled n=13), there are 256 possible combinations. Each combination is represented by an index (x-axis) and a RELSA score (y-axis, calculated as the average RELSA score of the analyzed animals (n=13) using the respective indexed variables/combinations). This RELSA score was analyzed separately for four different experimental time points: (**a**) post-op day, (**b**) day 1, (**c**) day 4, and (**d**) day 7. The top 5 and worst 5 combinations are highlighted purple or red, respectively.

#### Figure S4

##### S4a CLP nonsurvivors

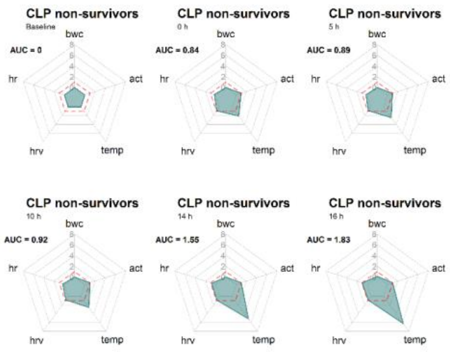


##### S4b CLP survivors

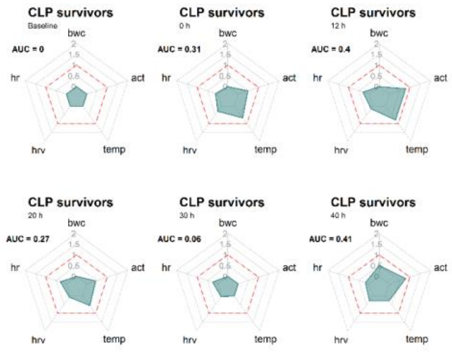


##### S4c CLP sham-operation

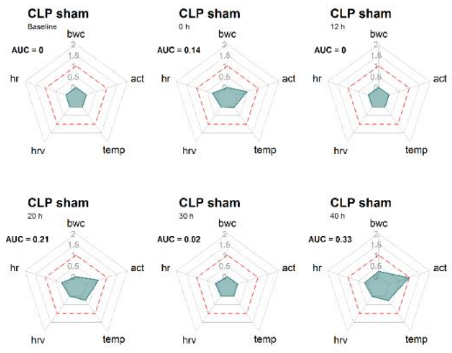


##### S4d DSS colitis

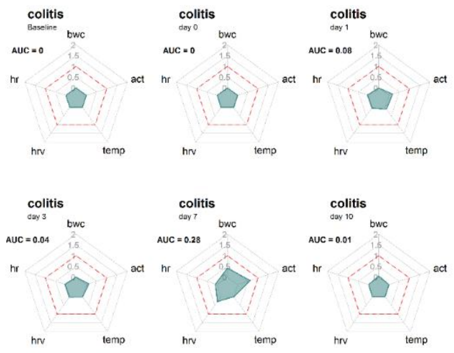


##### S4e DSS colitis + stress

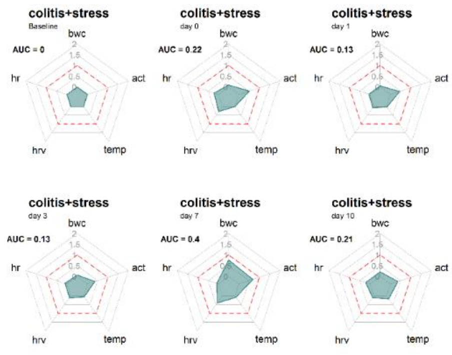


##### S4f DSS colitis control

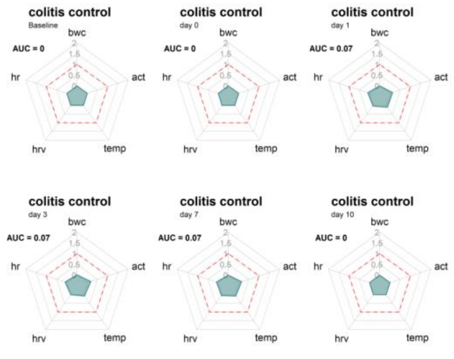


**Figure S4.** Radar charts of the RELSA weights ($R_w$) in the respective subgroups of the CLP (**a**-**c,** nonsurvivors, survivors, and sham) and DSS studies (**d**-**f**, colitis, colitis+stress and colitis control). For severity assessment and study comparisons of RELSA values, five variables were chosen that were available in all compared studies: body weight change (bwc), heart rate (hr), heart rate variability (hrv), body core temperature (temp) and activity (act). Depending on the study design, the $R_w$ were followed over different time points. The individual $R_w$ distributions reveal variables with strong or weak contributions to the RELSA score. The RELSA reference level maximum of 1 is symbolized with a red dashed line (RELSA=1). The area under the curve (AUC) value reports the standardized fraction of the RELSA reference level area (red dashed line) covered by the turquoise $R_w$ area (AUC=1 means 100% area coverage in all variables).

#### Tables

**S5.** Example of a table in the RELSA input format.

| | id | treatment | condition | day | bwc | mgs | bur2h | burON | hr | hrv | temp | act |
|---|---|---|---|---|---|---|---|---|---|---|---|---|
| 1 | Ca_001 | Transmitter | Carprofen | -1 | 100.00 | 0.10 | 56.40 | 71.94 | 453.56 | 12.43 | 36.61 | 1064.84 |
| 2 | Ca_001 | Transmitter | Carprofen | 0 | 87.50 | 0.11 | 14.92 | 44.69 | 660.59 | 2.05 | 36.18 | 125.54 |
| 3 | Ca_001 | Transmitter | Carprofen | 1 | 90.76 | 0.19 | 51.07 | 97.64 | 584.92 | 4.84 | 37.50 | 407.34 |
| 4 | Ca_001 | Transmitter | Carprofen | 2 | 92.93 | 0.17 | 42.28 | 99.01 | 538.04 | 6.77 | 37.14 | 351.64 |
| 5 | Ca_001 | Transmitter | Carprofen | 3 | 94.02 | NA | NA | NA | 482.16 | 7.07 | 36.71 | 286.28 |
| 6 | Ca_001 | Transmitter | Carprofen | 4 | 94.02 | 0.07 | 54.90 | 69.16 | 525.52 | 7.13 | 37.08 | 387.30 |
| 7 | Ca_001 | Transmitter | Carprofen | 5 | 96.74 | NA | NA | NA | 478.90 | 7.17 | 36.64 | 897.80 |
| 8 | Ca_001 | Transmitter | Carprofen | 6 | 97.83 | 0.14 | NA | NA | NA | NA | NA | NA |
| 9 | Ca_001 | Transmitter | Carprofen | 7 | 98.91 | NA | 45.42 | 52.73 | 472.28 | 7.69 | 36.32 | 707.06 |
| 10 | Ca_001 | Transmitter | Carprofen | 8 | 98.91 | NA | NA | NA | NA | NA | NA | NA |
| … | | | | | | | | | | | | |

**S6.** Overview of the animal numbers in the analyzed studies and subgroups.

| Mouse model | Strain | Sex | n |
|---|---|---|---|
| TM implantation metamizole | C57BL/6J | ♀ | 6 |
| TM implantation carprofen | C57BL/6J | ♀ | 7 |
| sham surgery metamizole | C57BL/6J | ♀ | 8 |
| sham surgery carprofen | C57BL/6J | ♀ | 7 |
| CLP | C57BL/6N | ♂ | 4 |
| CLP sham | C57BL/6N | ♂ | 3 |
| DSS colitis | C57BL/6J | ♀ | 8 |
| DSS colitis+restraint stress | C57BL/6J | ♀ | 7 |
| Colitis control | C57BL/6J | ♀ | 8 |
